# Young Infants Process Prediction Errors at the Theta Rhythm

**DOI:** 10.1101/2020.07.30.226902

**Authors:** Moritz Köster, Miriam Langeloh, Christine Michel, Stefanie Hoehl

## Abstract

Examining how young infants respond to unexpected events is key to our understanding of their emerging concepts about the world around them. From a predictive processing perspective, it is intriguing to investigate how the infant brain responds to unexpected events (i.e., prediction errors), because they require infants to refine their predictions about the environment. Here, to better understand prediction error processes in the infant brain, we presented 9-month-olds (*N* = 36) a variety of physical and social events with unexpected versus expected outcomes, while recording their electroencephalogram (EEG). We found a pronounced response in the ongoing 4 – 5 Hz theta rhythm for the processing of unexpected (in contrast to expected) events, for a prolonged time window (2 s) and across all scalp-recorded electrodes. The condition difference in the theta rhythm was not related to the condition difference in infants’ event-related activity to unexpected (versus expected) events in the negative central (Nc) component (.4 – .6 s), a component, which is commonly analyzed in infant violation of expectation studies using EEG. These findings constitute critical evidence that the theta rhythm is involved in the processing of prediction errors from very early in human brain development. We discuss how the theta rhythm may support infants’ refinement of basic concepts about the physical and social environment.

---

From early on, human infants develop basic concepts about their physical and social environment (Spelke and Kinzler, 2007). This includes a basic understanding of numbers (Wynn, 1992), the properties of objects (Baillargeon et al., 1985; Spelke et al., 1992), and others’ actions (Gergely et al., 2002; Reid et al., 2009). Our understanding of infants’ early concepts about their environment is based, to a large extent, on violation of expectation (VOE) paradigms. In VOE paradigms infants are shown unexpected events, which violate their basic concepts, in contrast to expected events. For example, infants are shown a change in the number of objects behind an occluder (Wynn, 1992), a ball falling through a table (Spelke et al., 1992), or an unusual human action (Reid et al., 2009). These unexpected events (in contrast to expected events) commonly increase infants’ attention. This is, for example, indicated by longer looking times. Unexpected events also motivate infants to learn about their environment, indexed by an increased exploration and hypothesis testing of objects which behaved unexpectedly (Stahl and Feigenson, 2015).

From a predictive processing perspective on infant brain development (Köster et al., 2020), VOE paradigms test infants’ basic predictions about physical and social events in their environment. To optimize their predictions and to reduce uncertainties, infants are thought to explore and integrate novel and unexpected information to reduce prediction errors in the long run (Köster et al., 2020; cf. Clark, 2013; Friston, 2011; Schubotz, 2015), which is reflected in their longer looking times or brain responses (please note that this basic idea also corresponds to classical conceptions of the response to VOE events; Sokolov, 1990). Yet, infants’ brain response to unexpected events is not fully understood. Based on the consideration that events that violate infants’ basic expectations elicit a prediction error and require infants to refine their predictions (Köster et al., 2020), the VOE response is highly interesting to better understand neural brain dynamics involved in prediction error processing in the infant brain.

Infants’ neural processing of unexpected events has formerly been investigated in terms of evoked neural responses (i.e., event-related potentials; ERPs) in the scalp-recorded electroencephalogram (EEG). This research has centered around the negative component (Nc), which emerges around 400-600 ms after stimulus onset at central recording sites, and which has been associated with attention processes (for a review, see Reynolds, 2015). However, unexpected events (in contrast to expected events) have been associated with an increased Nc (Kayhan et al., 2019; Langeloh et al., 2020; Reynolds and Richards, 2005; Webb et al., 2005) as well as a reduced Nc (Kaduk et al., 2016; Reid et al., 2009). Consequently, the neural mechanisms reflected in the Nc are not fully understood. Former studies have also investigated the spectral properties of the Nc component and linked this component to an increase in 1 – 10 Hz activity in infants and adults (Berger et al., 2006) or the 4 – 7 Hz activity for toddlers and adults (Conejero et al., 2018).

However, in the adult literature, most studies investigate the ongoing neural oscillatory activity (i.e., the oscillatory power within each trial, rather than the spectral characteristics of the ERP). It is critical to understand ongoing neural dynamics because they are fundamentally different from evoked oscillatory responses (Tallon-baudry and Bertrand, 1999) and their role in mnemonic processes, and the integration of novel information into existing representations in particular, has been well characterized in the adult brain (Friese et al., 2013; Hanslmayr et al., 2009; Klimesch et al., 1997; Köster et al., 2018; Osipova et al., 2006).

The 3 – 8 Hz theta rhythm plays an essential role in prediction error processing (Cavanagh and Frank, 2014) as well as learning processes in adults (Friese et al., 2013; Köster et al., 2018), children (Köster et al., 2017), and infants (Begus et al., 2015; Begus and Bonawitz, 2020). In a recent study, infants’ neural oscillatory dynamics were rhythmically entrained at 4 Hz, and the presentation of unexpected events led to a specific increase in the entrained 4 Hz, which was not the case for the entrained 6 Hz activity (Köster et al., 2019). However, it has not been investigated how the ongoing oscillatory activity responds to unexpected events in the infant brain and, specifically, whether the ongoing ∼ 4 Hz theta rhythm marks infants’ processing of prediction errors.

Here, we tested infants’ neural processing of prediction errors. We presented them with a series of different physical and social events with expected versus unexpected outcomes across various domains (see Figure 1), while recording their EEG. In particular, we used four different stimulus categories representing well-established paradigms from the VOE literature (testing infants’ concepts of action, solidity, cohesion, and number; see Figure S1 for the full stimulus set) to obtain a more generalized prediction error response, independent from a specific knowledge domain. Because of its pivotal role in prediction error processing and learning in adults, we expected a higher ongoing 4 Hz theta response for unexpected versus expected events. Furthermore, we expected a differential Nc response (400 – 600 ms, at central electrodes) between expected versus unexpected events and explored the associations between the Nc (and its spectral characteristics; cf. Berger et al., 2006) with the ongoing theta response.

**Figure 1.**
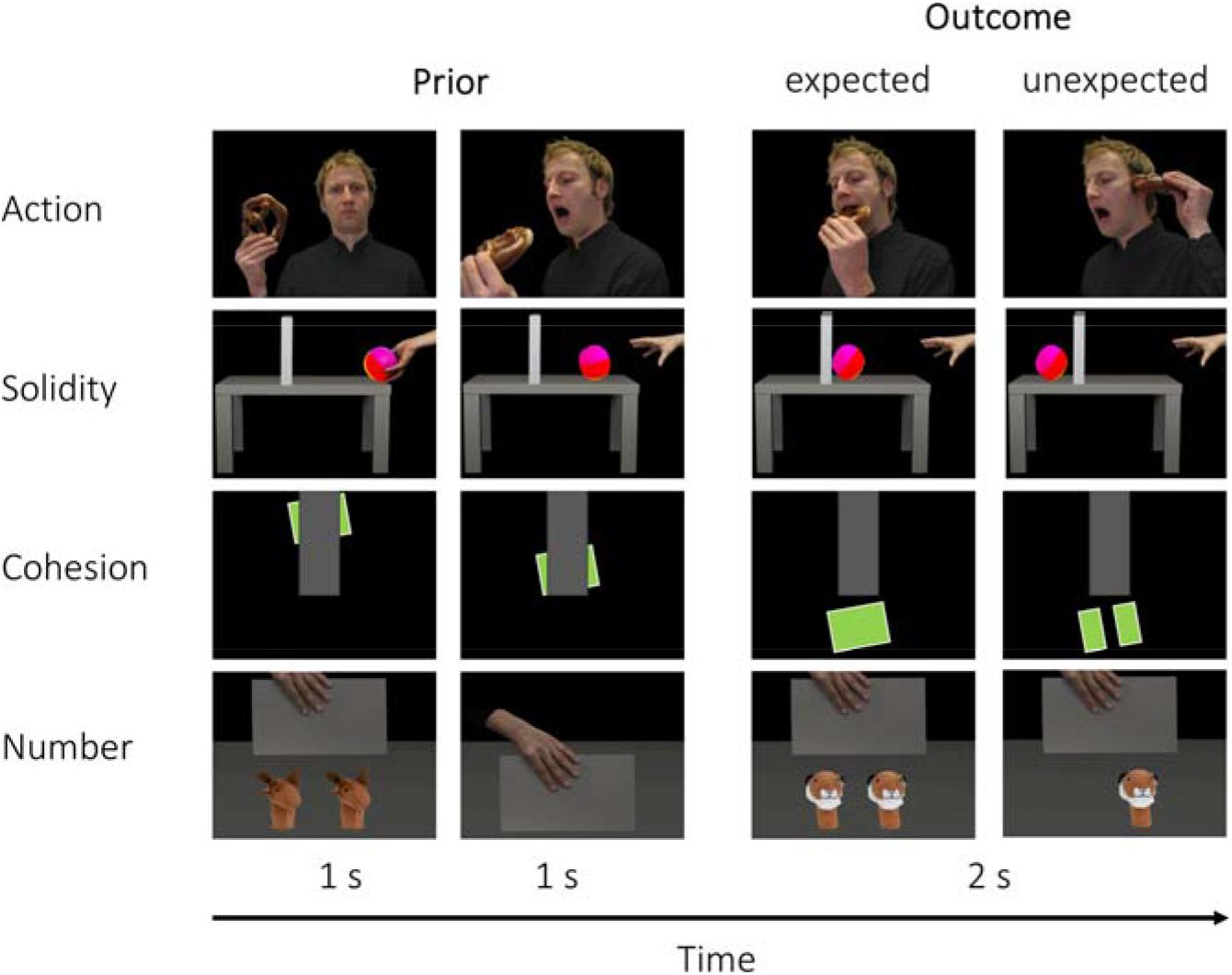
Examples of the violation of expectation events presented to participants. Infants saw the events of four basic knowledge domains (action, solidity, cohesion, and number). In each trial, the first two pictures initiated an event (prior; 1 s each) and the third picture showed the outcome (2 s), which could be expected or unexpected. (Please note that the outcome picture was shown for 5 s but we included all trials, in which infants watched for 2 s; see Stimuli and Procedure.)

## Materials and Methods

### Participants

The final sample consisted of 36 9-month-old infants (17 girls, *M* = 9.7 months, *SD* = 0.5 months). Participants were healthy full-term infants, from Leipzig, Germany. Informed written consent was obtained from each participant’s parent before the experiment and the experimental procedure was approved by the local ethics committee. Thirteen additional infants were tested but excluded from the final sample, due to fussiness (*n* = 2) or because fewer than 10 artifact-free trials remained in each condition (*n* = 11). This attrition rate is rather low for visual EEG studies with infants (Stets et al., 2012).

We selected this age group, because previous studies indicated VOE responses for the domains applied here, by the age of 9 months or even earlier (Reid et al., 2009; Spelke et al., 1992; Wynn, 1992). The sample size was based on a former study with a very similar study design (Köster et al., 2019) and ultimately determined by the number of families with infants in the targeted age-range, which were available in the period of the data assessment.

### Stimuli and Procedure

Stimuli were based on four classical VOE paradigms for the four core knowledge domains action, number, solidity, and cohesion, with four different stimulus types (variations) each, resulting in 16 different stimuli, which could be presented with an expected or unexpected outcome (see Figure S1, for the complete stimulus set). Each sequence consisted of three static images which, shown in sequence, depicted a scenario with a clearly predictable outcome.

In a within-subjects design, each of the 16 sequences was presented two times in each condition (expected or unexpected). This resulted in a total of 64 distinct trials, presented in 16 blocks. The order of the core knowledge domains, outcomes and the specific stimulus variations (four in each domain) were counterbalanced between blocks and across infants. We decided to present a high diversity of stimulus types from different domains to reduce transfer effects and keep infants’ attention high throughout the experiment. It should be noted that infants may get used to the stimuli and that this may reduce their surprise for unexpected outcomes over time. However, this would reduce, but not increase, the difference in the neural activity between expected and unexpected events.

Every trial began with an attention getter (a yellow duck with a sound, 1 s), followed by a black screen (variable duration of .5 – .7 s) and the three stimulus pictures. The first two pictures showed the initiation of an event or action (presented for 1 s each), followed by the picture presenting the expected or the unexpected outcome (see Figure 1). The final picture was presented for 5 s, for an accompanying eye-tracking study. Specifically, while we initially planned to assess and compare both infants’ gaze behavior and EEG response, the concurrent recording (EEG and eye-tracking) only worked for a limited number of infants and trials. Therefore, a match between the two measures was not feasible and we decided to collect more eye-tracking data in an independent sample, as a separate study. For the present study and analyses, we included all trials in which infants looked at the screen for at least 2 s of the final picture, coded from video (see below). The stimuli showing the outcome, namely the expected or unexpected outcome, were counterbalanced in case of the cohesion and the number stimuli (i.e., in the cohesion sequences outcome stimuli showed connected or unconnected objects and for number sequences the outcome showed one or two objects) and were matched in terms of luminance and contrast in case of the action and solidity stimuli (all *p* > .30). Stimuli were presented via Psychtoolbox (version 0.20170103) in Matlab (version 9.1). The full stimulus set of the original stimuli can be downloaded from the supplemental material of Köster et al. (2019).

Infants sat on their parent’s lap at a viewing distance of about 60 cm from the stimulus monitor. Sequences were presented at the center of a 17-inch CRT screen at a visual angle of approximately 15.0° × 15.0° for the focal event. We presented all 64 trials, but the session ended earlier when the infant no longer attended to the screen. On average, we presented *M* = 62.2, *SD* = 4.3 trials to the infants. A video-recording of the infant was used to exclude trials in which infants did not watch the first 4 s of a trial. Infants’ gaze behavior was coded offline, by a well-trained coder. We did not conduct a reliability coding, because the applied coding method has proven to be highly reliable. That is, in a former study with the same coding method we achieved an almost perfect agreement ICC= .979 (Köster et al., 2019).

### Electroencephalogram (EEG)

#### Apparatus

The EEG was recorded continuously with 30 Ag/AgCl ring electrodes from 30 scalp locations of the 10-20-system in a shielded cabin. Data were recorded with a Twente Medical Systems 32-channel REFA amplifier at a sampling rate of 500 Hz. Horizontal and vertical electrooculograms were recorded bipolarly, for the detection of eye-movements. The vertex (Cz) served as an online reference. Impedances were controlled at the beginning of the experiment, aiming for impedances below 10 kΩ.

#### Preprocessing

EEG data were preprocessed and analyzed in MATLAB (Version R2017b), using eeglab functions (Version v2019.1) and custom made scripts, applied in several former publications (e.g., Friese et al, 2013; Köster et al., 2018). EEG signals were band-pass filtered from 0.2 Hz to 110 Hz (using FIR filters) and segmented into epochs from -1.5 to 3 s, around to the onset of the outcome picture. Trials in which infants did not watch the complete 4 s sequence (2 s during the initiation of the event and 2 s of the outcome picture) were excluded from the analyses (number of trials removed: *M* = 23.1, *SD* = 7.1). Furthermore, noisy trials were identified visually and discarded (number of trials removed: *M* = 6.9, *SD* = 4.3) and up to three noisy scalp electrodes were interpolated based on spherical information. Thereafter, eye-blinks and muscle artifacts were detected using an independent component procedure (ICA; runica algorithm) and removed after visual inspection. To avoid any bias in the ICA removal, the ICAs were determined and removed across the whole data set, including all experimental conditions (both frequencies, both outcome conditions, all stimulus categories). Prior to the analyses, the EEG was re-referenced to the average of the scalp electrodes (Fz, F3, F4, F7, F8, FC5, FC6, Cz, C3, C4, T7, T8, CP5, CP6, Pz, P3, P4, P7, P8, Oz, O1, O2). Note that we only included scalp electrodes for the referencing, because mastoids and eye-channels should not contain brain signals (cf. Köster et al., 2019; Köster et al., 2020). Infants with a minimum of 10 artifact-free trials in each condition were included in the statistical analyses. Twenty-two to 52 trials (*M* = 32.2, *SD* = 7.3) remained for the infants in the final sample across conditions, with no significant differences in the number of trials between conditions (expected, unexpected), *t*(35) = 0.63 *p* = .530. We also plotted the data split by conditions, on subsamples with at least one trial for both the expected and the unexpected outcome condition. The respective size of subsamples and number of trials were action: *n* = 35, *M* = 10.3, *SD* = 3.2, solidity: *n* = 35, *M* = 6.9, *SD* = 2.7, cohesion: *n* = 32, *M* = 6.1, *SD* = 3.6, and number: *n* = 36, *M* = 8.5, *SD* = 3.0.

#### ERP Analysis

For the analyses of event-related potentials (ERPs), we averaged the neural activity, separately for the trials of both conditions (expected, unexpected). We focused on the NC as a classical component associated with infants’ processing of expected versus unexpected events (Reynolds, 2015). Specifically, we averaged the ERPs across central electrodes (Cz, C3, C4), and between 400 – 600 ms, with regard to a -100 – 0 ms baseline. We chose a baseline just before the onset of the outcome picture, because it was part of the picture sequence, and each picture elicited a neural response, and this response (4 – 5 Hz and ERP) decayed towards the beginning of the next stimulus (see Figure S2). The ERP power was averaged for each participant and condition and the power between expected and unexpected trials was then contrasted by means of a dependent *t*-test. We band-pass filtered the ERPs from 0.2 – 30 Hz for displaying purposes.

#### Spectral Analysis

To obtain the trial-wise spectral activity elicited by the outcome pictures we subjected each trial to a complex Morlet’s wavelets analysis (Morlet parameter *m* = 7, at a resolution of 0.5 Hz). We then averaged the spectral power across trials, separately for conditions (expected, unexpected). We focused on the frequencies between 2 and 15 Hz across the whole analyzed time window 0 – 2000 ms, with regard to the -100 – 0 ms baseline. This was to obtain the neural response to the outcome pictures and also to make the results directly comparable to the ERP analysis in this and former studies. We did not analyze higher frequencies due to muscle and ocular artifacts in the infant EEG (e.g., Köster, 2016).

Because this was the first study to look at the trial-wise neural oscillatory response to a series of unexpected versus expected events (i.e., not only including neural dynamics tightly locked to the stimulus onset, reflected in the ERP; cf. Berger et al., 2006), in a first step, we looked at the grand mean spectral activity, separated by conditions (unexpected, expected), and the difference between both conditions (unexpected -expected). Conservatively and because we did not have a specific hypothesis about the topography or temporal evolution of the theta rhythm across all domains, we analyzed the neural oscillatory activity averaged across the whole time-range of the outcome stimulus (0 – 2000 ms) and all scalp-recorded electrodes (Fz, F3, F4, F7, F8, FC5, FC6, Cz, C3, C4, T7, T8, CP5, CP6, Pz, P3, P4, P7, P8, Oz, O1, O2). This relies on our former studies, where we entered into the analysis (and found a theta effects across) the whole time window of analysis and across all scalp recorded electrodes (Köster et al., 2017; Köster et al., 2018; Köster, Martens & Gruber, 2019). Note that this method is highly conservative, since no specific time-window nor specific electrodes are selected. For this reason, no multiple comparison correction for the selection of time and electrode space was applicable. Our initial proposal was that the 4 Hz theta rhythm would be higher for unexpected, compared to expected events (Köster et al., 2019). However, the analysis for the 4 Hz frequency (technically 3.9 Hz) only reached the threshold of a one-sided significance (*t*[35] = 1.29, *p* = .102, *d* = 0.24; one-sided *p* = .051), while the adjacent frequencies (4.4 and 4.9 Hz) differed significantly between conditions (*t*[35] = 2.22, *p* = .017, *d* = 0.45, and, *t*[35] = 2.60, *p* = .007, *d* = 0.49). For this reason, and because the present study was the first to conduct this specific analysis for infants VOE response in the EEG, we included all 3 frequency bands (3.9, 4.4, and 4.9 Hz, hereafter 4 – 5 Hz) in the main analyses.

## Results

Infants’ event-related responses upon the onset of the outcome picture revealed a clear Nc component between .4 – .6 s over central electrodes. The Nc was more pronounced for expected in contrast to unexpected events, *t*(35) = -2.87, *p* = .007, *d* = 0.60 (Figure 2).

**Figure 2.**
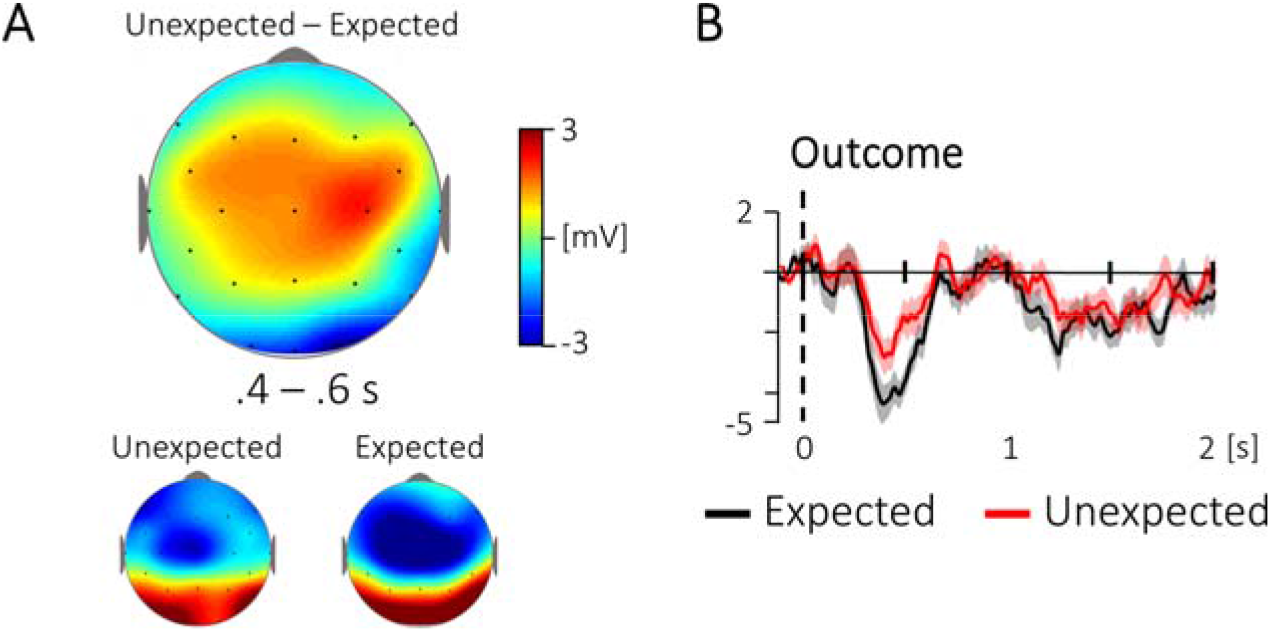
The topography and time course of the Nc for the outcome pictures. (A) The upper topography shows the difference between unexpected and expected events for .4 – .6 s, in contrast to a -100 – 0 ms baseline. The lower topographies show the activity for expected and unexpected events, separately. (B) The corresponding time course at central electrodes (Cz, C3, C4), with a significant difference between .4 – .6 s, *p* = .007.

Furthermore, across all scalp recorded electrodes and the whole 0 – 2 s time window, we observed an increase in neural oscillatory activity in the 4 – 6 Hz range for unexpected events (Figure 3A, upper panel) and an increase at 6 Hz for expected events (Figure 3A, middle panel). We verified this observation with corresponding t-tests, *t*(35) = 4.38, *p* < .001, *d* = 0.73, and, *t*(35) = 4.57, *p* < .001, *d* = 0.76. This resulted in higher 4 – 5 Hz activity for unexpected compared to expected events across all scalp-recorded electrodes and throughout the whole 0 – 2000 s time-window, *t*(35) = -2.34, *p* = .025, *d* = 0.37 (Figure 3B and C).

**Figure 3.**
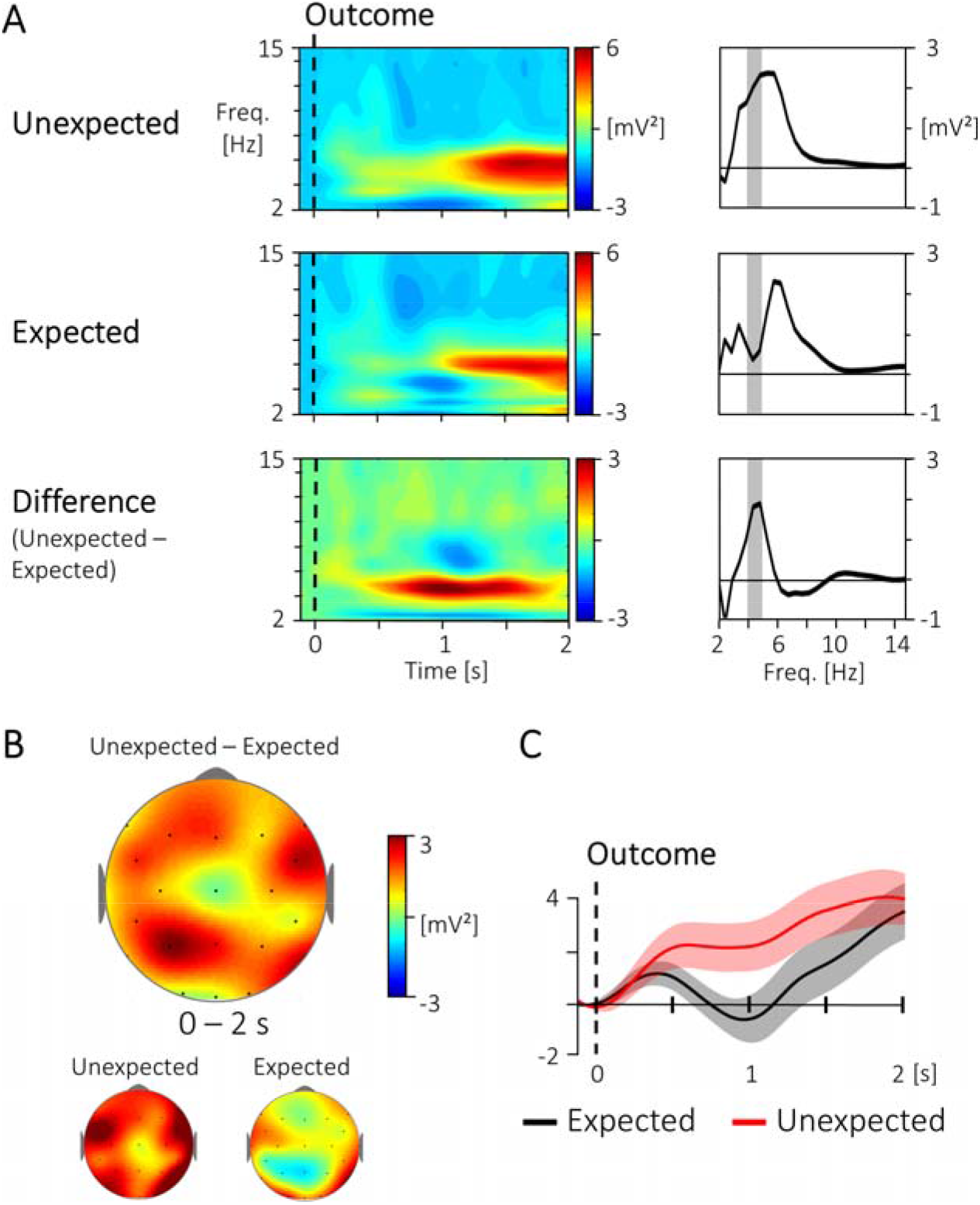
The grand mean spectral characteristics for unexpected versus expected events for the outcome picture. (A) The left panels show the time-frequency response across all scalp-recorded electrodes for unexpected and expected events and the difference (unexpected – expected), with regard to a -.1 – 0 s baseline. The right panels show the frequency response between 2 and 15 Hz, averaged over time. The gray area indicates the 4 – 5 Hz range. (B) The upper topography shows the unexpected -expected difference in 4 – 5 Hz activity across the whole time window of analysis (0 – 2 s, baseline: -.1 – 0 s). The lower topographies show the activity for expected and unexpected events, separately. (C) The corresponding time course for the 4 – 5 Hz response across all scalp-electrodes and the whole 0 – 2 s time window shows a significant difference between unexpected versus expected events, *p* = .025.

To investigate the relation between the effects which we found in the ERP and the ongoing 4 – 5 Hz theta activity, we tested the spectral characteristics of the evoked oscillatory activity (i.e., by applying a wavelet transform to the ERP) and its relation to the ongoing oscillatory activity at central electrodes (Cz, C3, C4) between .4 – .6 s (Figure S3). We did not find a significant condition difference in the evoked activity, *t*(35) = 1.57, *p* = .126, *d* = 0.34, nor the ongoing activity, *t*(35) = -1.26, *p* = .218, *d* = 0.29, at the central electrodes. The condition differences (unexpected -expected), between .4 – .6 s at central electrodes, were also not correlated between the ongoing and the evoked response, neither for the difference in the actual ERP, *r* = -.02, *p* = .919, nor its spectral characteristics, *r* = .23, *p* = .169. This indicates that the ongoing theta response captures a different characteristic of the EEG signal than reflected in the ERP.

Although the present study was designed to investigate infants’ processing of prediction error in general, across domains, to get an impression about the consistency of the differences in the central Nc and the 4 – 5 Hz activity, we plotted the data split by domains (action, solidity, continuity, number). Descriptively, the overall time course of the ERP and the 4 – 5 Hz effect was somewhat consistent across conditions, however, the condition differences (Figure 1 and 2) seem to be driven, to a large degree, by the stimuli of the action and the number domain (see Figure S4 and S5). Interestingly, the peak in the unexpected – expected difference was in the 4 – 5 Hz range across all four domains (Figure S5 A). Critically, we did not test domain-specific differences statistically due to the low trial numbers within each domain.

## Discussion

Our results show a clear increase in the ongoing 4 – 5 Hz power in response to unexpected events, in contrast to expected events, across all scalp-recorded electrodes and the whole time window of 2 s, after the onset of unexpected outcome pictures. Thus, the theta rhythm was substantially increased for the processing of prediction errors in the infant brain. In the ERP response we found a stronger Nc for expected events, in contrast to unexpected events, at central electrodes.

A direct comparison at central electrodes revealed that the effects of the ongoing theta response do not reflect the condition differences in the ERP nor in the spectral characteristics of the ERP (cf. Tallon-baudry and Bertrand, 1999). In fact, both condition differences pointed into the opposite direction (unexpected < expected) than the difference in the ongoing theta activity (expected < unexpected). Thus, the ongoing theta response characterizes a distinct neural signature compared to the ERP analyses reported in former studies (Berger et al., 2006; Conejero et al., 2018; Kayhan et al., 2019; Langeloh et al., 2020; Reynolds and Richards, 2005; Webb et al., 2005).

The ongoing theta rhythm has been associated with learning processes in human adults (Friese et al., 2013; Hanslmayr et al., 2009; Klimesch et al., 1997; Köster et al., 2018; Osipova et al., 2006), children (Köster et al., 2017), and infants (Begus et al., 2015). Our findings highlight that the theta rhythm promotes the processing of novel, unexpected information, in the sense of prediction errors, already in early infancy. This is particularly interesting because the theta rhythm is usually associated with neural processes in prefrontal and medio-temporal structures, which are still immature in the infant brain (Gilmore et al., 2012). Besides its involvement in forming novel representations, the theta rhythm has long been associated with cognitive control processes in adults (Cavanagh and Frank, 2014; Hanslmayr et al., 2008) and children (Adam et al., 2020), and infants’ ongoing theta oscillations at 6 months were predictive for their cognitive ability at 9 months (Braithwaite et al., 2020). However, in adults the theta rhythm is mostly found in prefrontal brain regions and the present results and a former SSVEP study (Köster et al., 2019) indicate that the theta rhythm may be pronounced in parietal networks in the infant brain. While we do not want to overemphasize the topography in the EEG of infants and also did not test the location empirically (for the same reason), it may be that the theta rhythm plays a role in building up semantic knowledge structures in parietal cortical networks (cf. Köster et al., 2017). However, further studies are needed to more clearly characterize the topography and specific neural processes reflected in the infant theta rhythm.

We also identified differences between unexpected and expected events in the Nc, a classical visual ERP component associated with infants’ processing of unexpected events. As expected from former studies, the Nc and the condition difference was pronounced between 400 – 600 ms, and was specific to central electrodes (Cz, C3, C4). However, the condition difference pointed in the opposite direction than most (Kayhan et al., 2019; Langeloh et al., 2020; Reynolds and Richards, 2005; Webb et al., 2005), though not all (Kaduk et al., 2016; Reid et al., 2009), previously reported Nc effects (namely, the more common findings of a higher negativity for unexpected events). It is currently not clear, why unexpected events induce enhanced Nc amplitudes in some studies, but a decreased Nc compared to expected events in others. Because the amplitude of the Nc has been associated with the extent of attentional engagement with a visual stimulus (Reynolds, 2015; Reynolds and Richards, 2005), in our study infants’ initial orienting response may have been more pronounced for the more familiar and expected outcomes. This is in line with previous studies using partly similar stimuli (in particular the action events; Kaduk et al., 2016; Reid et al., 2009) and with the notion that infants show familiarity preferences (i.e., the preference for events consistent with their experience) when they are still in the process of building stable cognitive representations of their environment (Nordt et al., 2016).

While the present study is, to the best of our knowledge, the first to look at a generalized prediction error (i.e., VOE) response in the ongoing theta rhythm, we would also like to clearly indicate its limitations. To obtain a generalized response we presented stimuli from 4 different knowledge domains. Although we obtained reasonable trial numbers for the overall VOE response, this resulted in a low number of trials for each individual domain. Despite some descriptive similarities across those domains, there also seems to be considerable variation in the magnitude and topographies of the described effects (see Figures S5 and S6). However, more trials in each domain would be needed to resolve these empirically. Furthermore, we applied a conservative selection of electrodes (i.e., all electrodes) and the analyzed time window (i.e., the whole 2 s time window), but did not find a clear effect at the predicted 4 Hz frequency (cf. Köster et al., 2019). Instead, the condition difference was highest at the adjacent frequencies (4.5 and 5 Hz), and the selection of 4 – 5 Hz should therefore be confirmed in further studies. Finally, the selection of a short -1. – 0 s baseline was needed to test the changes in the ongoing theta rhythm within a continuous sequence of stimuli (and make it directly comparable to the ERP analyses), while the theta rhythm within the trial may have influenced this baseline. However, because we found increased theta activity and higher theta for unexpected versus expected events, this may have reduced, but could not explain, the critical difference (unexpected > expected).

Embedding the role of the theta rhythm in a broader theoretical framework, from animal models we know that the theta rhythm encodes predictions (i.e., such as the activation of future locations in a labyrinth; O’Keefe and Recce, 1993) and facilitates Hebbian learning (Tort et al., 2009). Based on these findings, the theta rhythm has been described as a neural code for the sequential representation and the integration of novel information into existing concepts (Lisman and Jensen, 2013). We would like to add to this that the theta rhythm may implement a computational mechanism that compresses real time events onto a faster neural time-scale. This may serve the function to advance with cognitive processes ahead of real time and to facilitate the integration of new events into existing networks. That is, speeding up real time events may enable the brain to predict future events and integrate novel events as they happen in real time (e.g., such as the rat, predicting where it will arrive next in the labyrinth). It is critical that neural events, which happen in real time, are activated at the neural level, in close temporal proximity, to allow long-term potentiation processes (Hebb, 1949; cf. Köster et al., 2018). Former studies have demonstrated that computational mechanisms reflected in the theta rhythm may be phylogenetically preserved in the mammalian lineage (Cavanagh and Frank, 2014; Lisman and Jensen, 2013). Here we report first evidence that the ongoing theta rhythm is present during the processing of unexpected events already from very early in human ontogeny.

To conclude, our findings make a strong case that the theta rhythm is present from very early in ontogeny, associated with the processing of prediction errors and, possibly, the refinement of the emerging concepts of the physical and social environment. This marks an important step towards a better understanding of the neural oscillatory dynamics that underlie infants’ brain development and their emerging models of the world around them.

## Acknowledgements

We would like to thank Carl Bartl und Ulrike Barth for their support with the data assessment and coding.

