## Supplementary material for "Young Infants Process Prediction Errors at the Theta Rhythm"

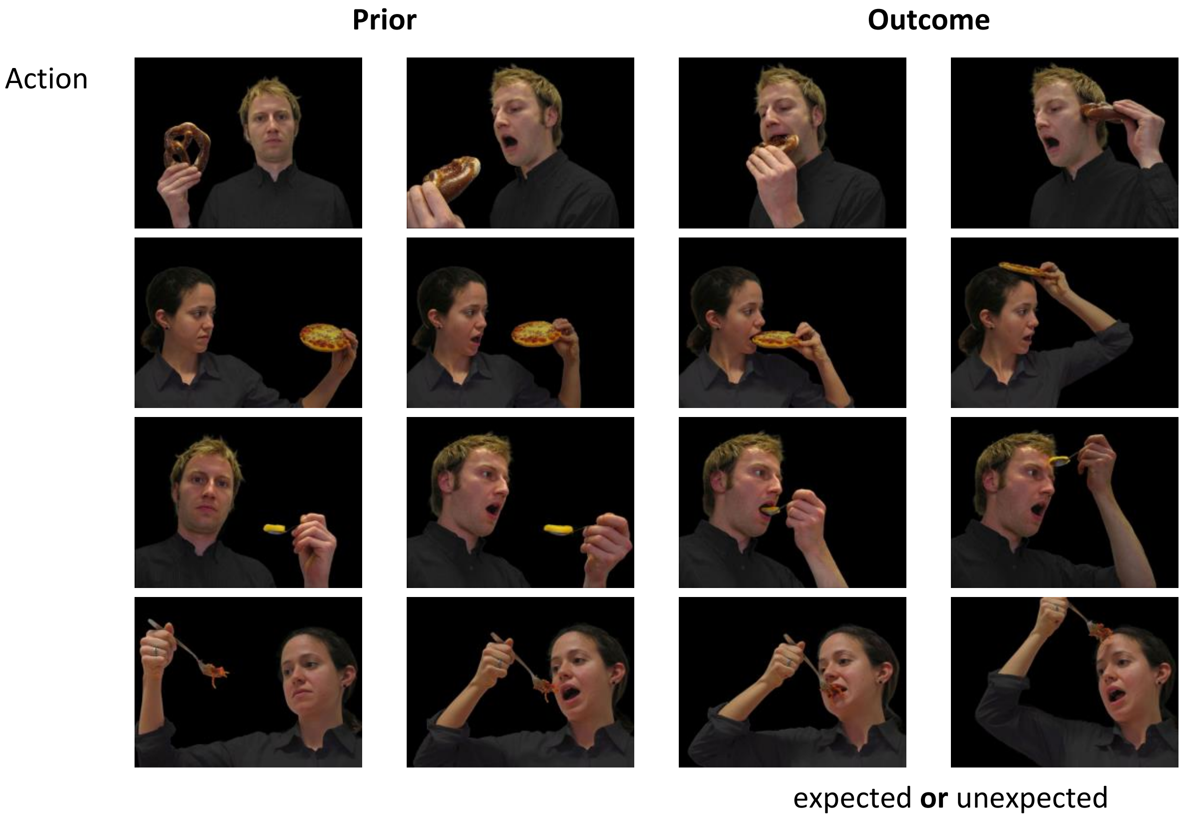
Prior
Outcome
Action
expected **or** unexpected

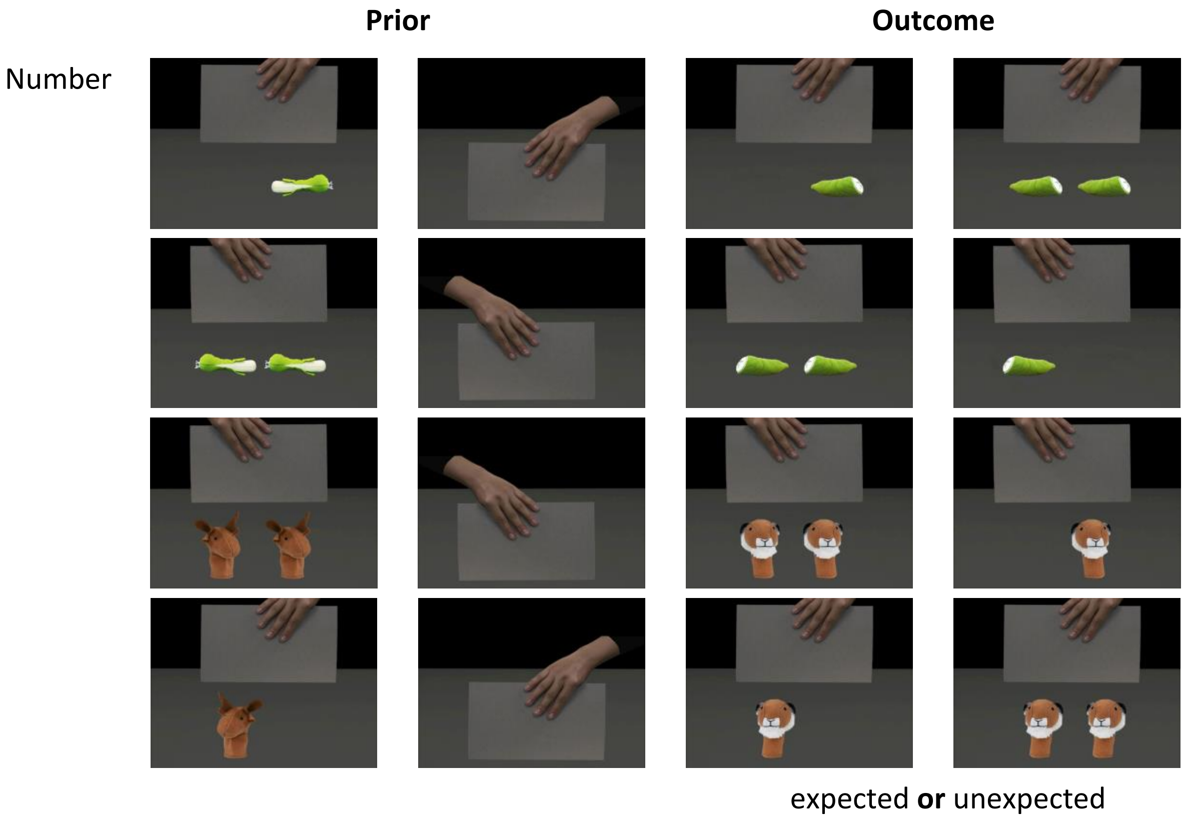
Prior
Outcome
Number
expected **or** unexpected

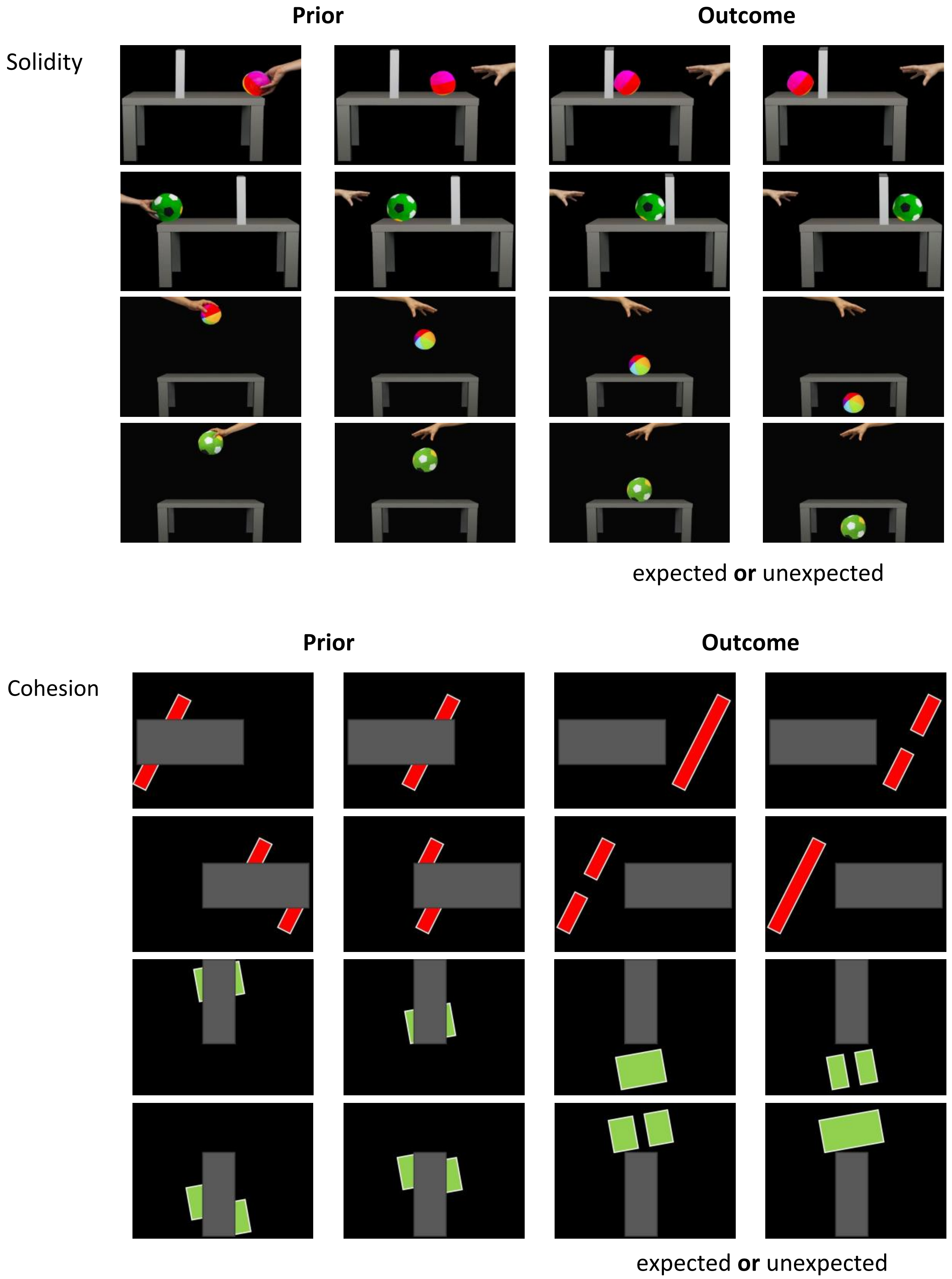


**Figure S1.** Stimulus sets. Complete stimulus sets used in the (A) action, (B) number, (C) solidity and (D) cohesion domain. Each trial consisted of three stimulus pictures. The first two pictures showed the initiation of an event or action (prior and baseline picture) followed by the third picture, presenting either an expected or an unexpected outcome.

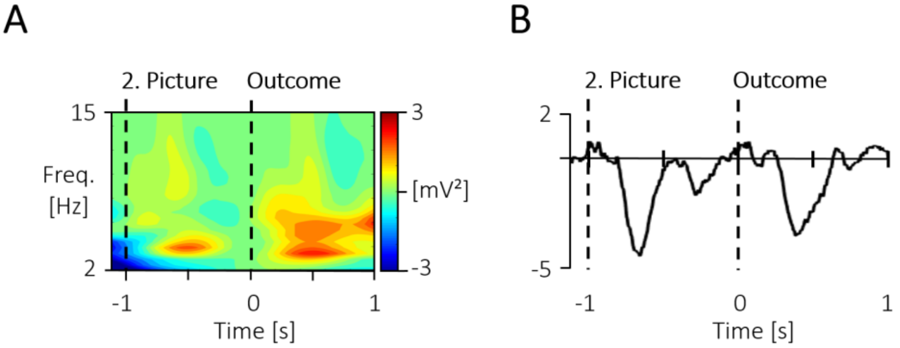


**Figure S2.** Display of the grand mean signals prior to the -100 to 0 baseline and outcome stimulus. (A) The time-frequency response across all scalp-recorded electrodes, as well as (B) the ERP at the central electrodes (Cz, C3, C4) across both conditions show a similar response to the 2. picture like the outcome picture. Thus a baseline just prior to the stimulus of interest was chosen, consistent for both analyses.

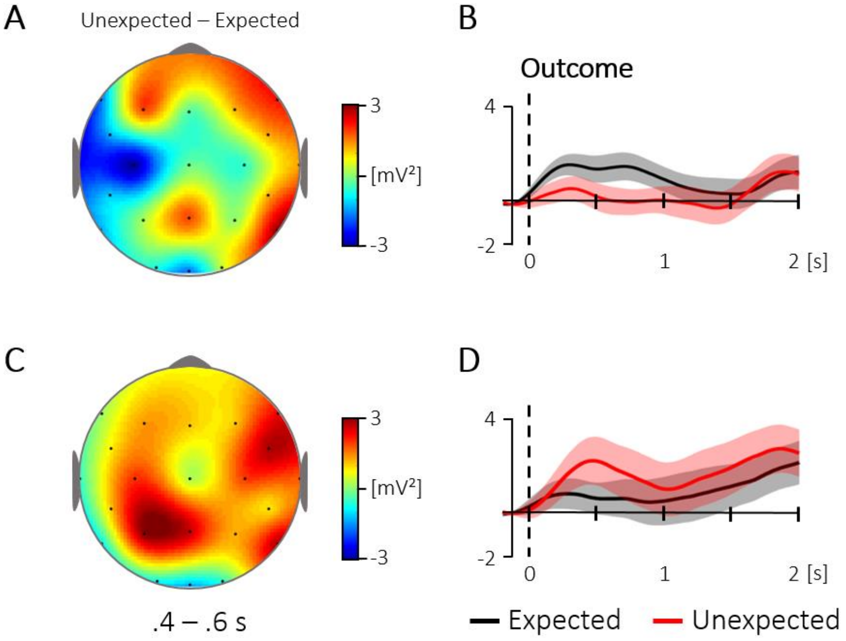


**Figure S3.** Comparison of the topography and time course for the 4 – 5 Hz activity for the evoked (A, B) and the ongoing (C, D) oscillatory response at central electrodes (400 – 600 ms; Cz, C3, C4; baseline: -100 – 0 ms). (A) Topography for the difference between unexpected and expected events in the evoked oscillatory response. (B) The corresponding time course at central electrodes, which did not reveal a significant difference between 400 – 600 ms, t(35) = 1.57, p = .126. (C) Topography for the same contrast in the ongoing oscillatory response and (D) the corresponding time course at central electrodes, which did likewise not reveal a significant difference between 400 – 600 ms, t(35) = -1.26, p = .218. The condition effects (unexpected - expected) were not correlated between the evoked and the ongoing response, $r$ = .23, $p$ = .169.

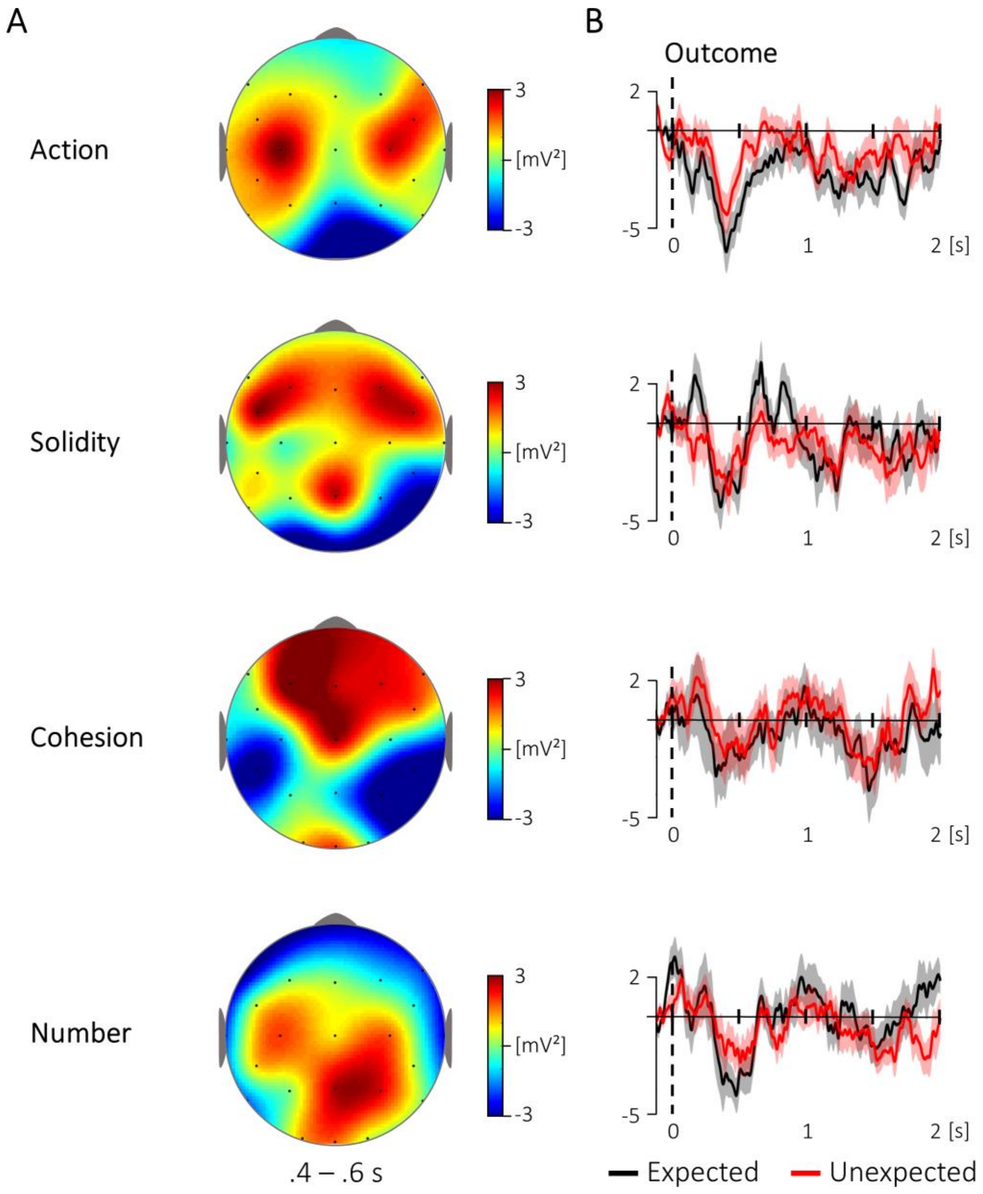


**Figure S4.** The topography and time course of the NC for the outcome pictures, split by the different knowledge domains. (A) The difference between unexpected and expected events for 400 – 600 ms, in contrast to a -100 – 0 ms baseline. (B) The corresponding time course at central electrodes (Cz, C3, C4). Note that these plots show different subsamples (see Materials and Methods for details).

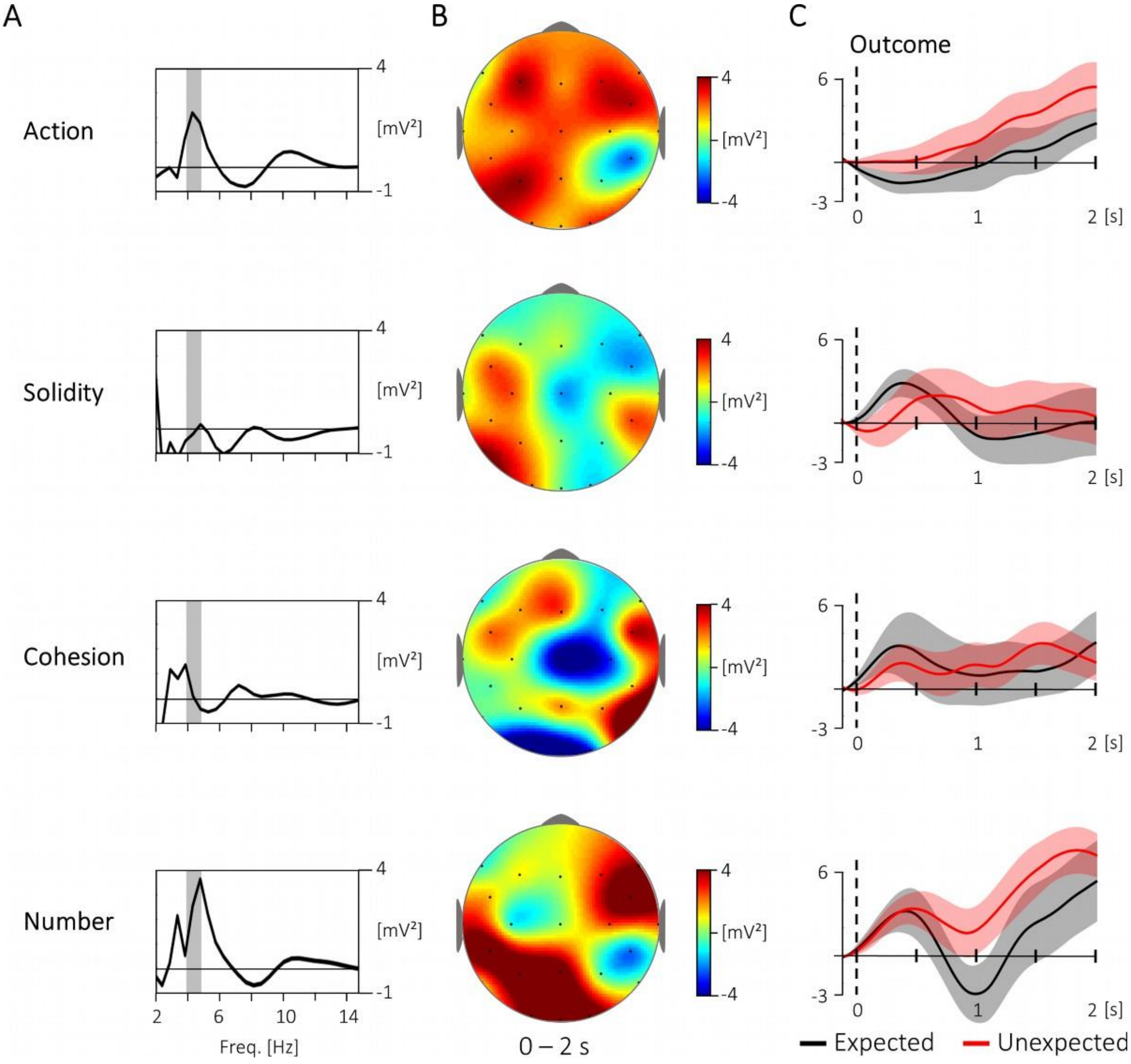


**Figure S5.** Spectral characteristics for unexpected versus expected outcome pictures, split by the different knowledge domains. (A) The frequency responses between 2 and 15 Hz, averaged across all electrodes and the whole time window (0 – 2000 ms, baseline: -100 – 0 ms). The gray area indicates the 4 – 5 Hz range. (B) The topography shows the unexpected - expected difference in 4 – 5 Hz activity across the whole time window (0 – 2000 ms, baseline: -100 – 0 ms). (C) The corresponding time course for the 4 – 5 Hz response across all scalp-electrodes and the whole 0 – 2000 ms time window. Note that these plots show different subsamples (see Materials and Methods for details).
